# CrossLabFit: A Novel Framework for Integrating Qualitative and Quantitative Data Across Multiple Labs for Model Calibration

**DOI:** 10.1101/2024.12.08.627398

**Authors:** Rodolfo Blanco-Rodriguez, Tanya A. Miura, Esteban Hernandez-Vargas

## Abstract

The integration of computational models with experimental data is a cornerstone for gaining insight into biomedical applications. However, parameter fitting procedures often require a vast availability and frequency of data that are challenging to obtain from a single source.

Here, we present a novel methodology “CrossLabFit” designed to integrate qualitative data from multiple laboratories, overcoming the constraints of single-lab data collection. Our approach harmonizes disparate qualitative assessments—ranging from different experimental labs to categorical observations—into a unified framework for parameter estimation. By using machine learning algorithms, these qualitative constraints are represented as dynamic “qualitative windows” that capture significant trends to which models must adhere. For numerical implementation, we developed a GPU-accelerated version of differential evolution to navigate in the cost function that integrated quantitative and qualitative data.

We validate our approach across a series of case studies, demonstrating significant improvements in model accuracy and parameter identifiability. This work opens a new paradigm for collaborative science, enabling a methodological road to combine and compare findings between studies to improve our understanding of biological systems and beyond.

## Introduction

Computational modeling is a fundamental endeavor that enables the exploration of elusive biological phenomena. Mathematical models have unknown parameters whose estimation is of paramount importance. Without accurate parameter values, the model predictive value is compromised [1]. Rigorous parameter estimation not only verifies the model’s efficacy but also refines our understanding of the biological phenomena in question [2].

Maximum likelihood estimators find values of the model parameters that give the observed data the highest probability [3]. This can be achieved by maximizing the likelihood profile or minimizing a cost function such as the residual sum of squares, where the best parameter set lies at the minimum of the cost function landscape. A computational technique employing profile likelihood was introduced by [4] to determine model parameters in ordinary differential equations based on experimental data. This method also enables the detection of structural and practical non-identifiability [5].

A common practice in the modeling community is reducing model complexity to tackle identifiability problems, however, this dramatically limits the holistic understanding of the biological problem [6]. Another approach is to integrate new data sets into the parameter fitting procedure [2]. Nevertheless, acquiring new data is a daunting and expensive endeavor, often constrained by resource limitations and experimental feasibility. Another common and debatable practice in the modeling community is bringing data sets from different research articles. Even when laboratories rely on available biological standards to allow for reproducible quantification [7], quantitative inconsistencies still arise.

Datasets in biology cannot be compared straightforwardly between laboratories in a quantitative form, and it is not necessarily because of the stochasticity of biology [8]. For example, cell quantification by flow cytometry data is dependent on cell collection and staining techniques [9], and quantification of specific mRNAs by quantitative PCR is dependent on reaction conditions. Data from these examples are also affected by instruments and user-defined settings [10].

Quantification of virus concentrations provides another example of the difficulty in deriving accurate quantitative values that can be directly compared across studies [11]. Knowing the concentration of virus stocks is essential for calculating accurate dosages for infection studies and measuring the effectiveness of antiviral compounds. Within studies, these values can be reported as relative concentrations compared to values from control samples, but they are rarely comparable between studies [12]. Differences between laboratories and even replicate assays within the same lab can result prevent direct comparison of data from plaque assays. These differences can include differences in cell line passage history, details of the assay protocol, cell density, and experimental error, all of which can affect the ability to visualize and accurately count plaques [13].

The use of qualitative data emerges as a potential approach to increase the accessibility of experimental data. The study in [14] proposed a methodology that combines qualitative and quantitative data. Their methodology hinges on constructing a unified objective function. Within this framework, a conventional quantitative term, expressed as the standard sum of squares over all data points, coexists with a qualitative term. This qualitative component incorporates observations in the form of inequality constraints, penalizing the cost function accordingly. This approach was implemented in the pyBioNetFit toolbox [15] and extended with Bayesian inference [16]. Another way to integrate data is by using monotonic relationships between the experimental measure and simulated model data, a concept known as optimal scaling introduced in [17, 18]. Optimal scaling was achieved by categorizing data and generating surrogate data while preserving the monotonicity observed in experimental data [19]. Integrating gradient-based methods with qualitative data significantly improves the accuracy and speed of parameter estimation [20, 21]. This approach has been implemented in the Python Parameter EStimation TOolbox (pyPESTO) [22].

Despite these approaches, qualitative behavior is grossly defined to upper or lower bounds, *e*.*g*., qualitative data are converted into inequality constraints imposed on the outputs of the model [14] or to categories [20]. Additionally, it is assumed that qualitative data come from the same lab group’s experiments. However, there is a wealth of untapped data sets in the scientific literature that can be merged to define temporal qualitative categories. For example, influenza dynamics is used to estimate parameters of the replication cycle and enable forecasting of future dynamics [23–26]. Nonetheless, interactions with different arms of the immune system would require additional data [27–30], which is very unlikely to be all from the same lab.

Here, we propose a new methodology based on machine learning to generate qualitative categories by merging experimental data from different laboratories. Based on the generated qualitative data, we apply a novel approach called “CrossLabFit”. In this approach, the cost function is divided into one fitting quantitative data and the other penalizing deviations from the unobserved variables. This approach allows us to limit the distributions in model parameters and limit model trajectories from those that are not realistic.

## Results

The main idea behind CrossLabFit is to model a data set of interest (Fig 1a) by minimizing a cost function *e*.*g*. residuals. As the data we want to explain is very likely limited to a few variables, we propose to integrate additional data sets from different labs and sources into dynamic domains where the model trajectories should reside, see Fig 1b). We refer to this dynamic domain as *qualitative windows*. These qualitative windows will delineate the search parameter space by penalizing the cost function on model trajectories that do not pass through a qualitative window (Fig 1c). Qualitative windows will refine the landscape of search for the global minimum while limiting model trajectories that have unrealistic biological behavior (Fig 1d). By using a test-bed model, we will show the efficacy of the integrative cost function next.

**Fig 1.**
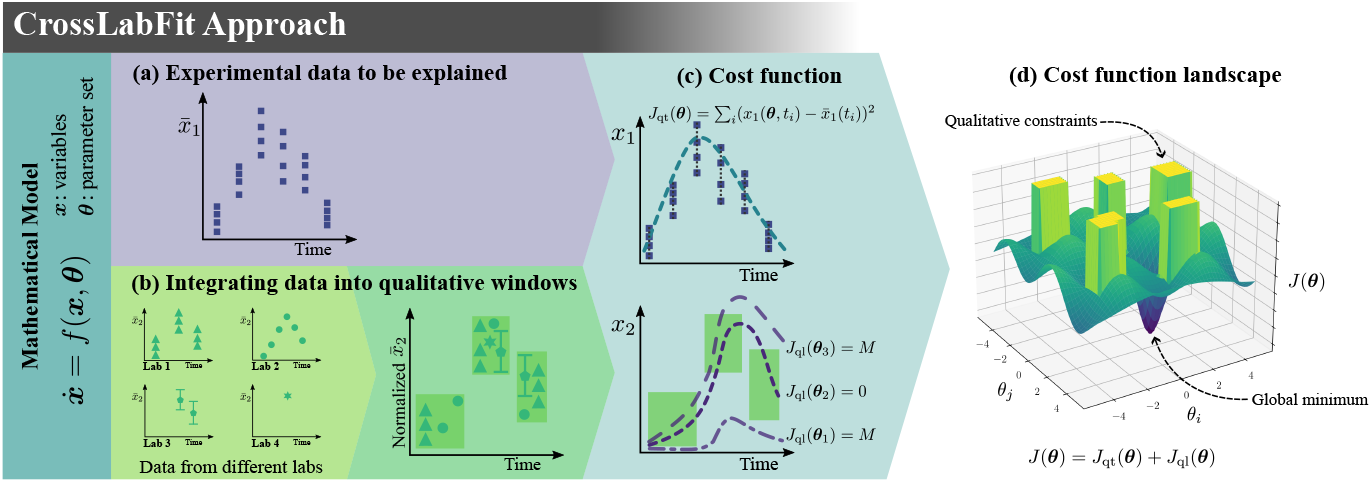
CrossLabFit approach. It is consider an experimental data to be explained (a). Additional information from various sources is collected to build qualitative window constraints (b). By incorporating these constraints into the cost function (c), we transform the cost function landscape to enhance the search for the global minimum (d).

### Integrative cost function

While CrossLabFit approach can be applied to any computational model, for presentation purpose, we will consider here a mathematical model expressed with Ordinary Differential Equations (ODEs) to study a biological problem in the following form

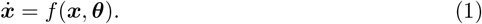

Model variables ***x*** are defined by equations that depend on these variables and a set of parameters ***θ*** = (*θ*_1_, *θ*_2_, …, *θ*_*n*_). We typically fit our model by finding the minimum of a cost function that measures the difference between our simulated variable *x*_*i*_ and the empirically observed data 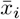.This cost function corresponds to a landscape in a hyperspace ℝ^*n*^, where *n* is the number of parameters in ***θ***.

The essence of our approach lies in the optimization routine, which requires the reformulation of the cost function to include qualitative data. The resulting cost function is a composite of quantitative and qualitative elements given by

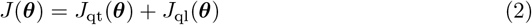

where *J*_qt_ and *J*_ql_ represent the cost functions corresponding to the quantitative and qualitative parts, respectively. For the quantitative part, we consider the Residual Sum of Squares (RSS) in the following form

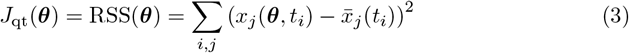

where *x*_*j*_ is the observed variable, 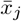 is the empirical data of the variable *j*, and *t* is the time.

We integrate the qualitative component of the cost function as penalties for cases where the model trajectories fall outside these qualitative windows. These constraints apply to variables not included in the quantitative fit. Each qualitative window encompasses a specific range of time and variable values, thereby defining the qualitative cost function as

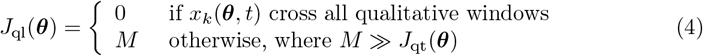

where *x*_*k*_ are the variables fitted to the qualitative data and *M* is a penalty factor much larger than *J*_qt_.

Fig 1d) illustrates the qualitative component of the cost function represented as rectangular barriers in the cost function landscape. These high barriers in the landscape eliminate model trajectories, preventing further exploration in those areas. In addition, these barriers not only speed up the search for the best parameter set but also exclude parameter sets that do not correspond to realistic biological behavior, improving parameter identifiability.

### Benchmark to evaluate performance

To illustrate the potential of this integrative cost function, we used three-species Lotka-Volterra models, each tailored to represent different interaction dynamics within a system. In this section, we present the results of the cycle Lotka-Volterra model, which is shown with its corresponding schematic interactions and equations in Fig 2; the other two variations are presented in Supplementary Material Fig S2 and Fig S3. This model represents a cyclic interaction among three species, with the third species influencing the first, creating a feedback loop. This model serves as a testbed to assess the efficacy of the CrossLabFit approach when qualitative data are synergized with quantitative data.

**Fig 2.**
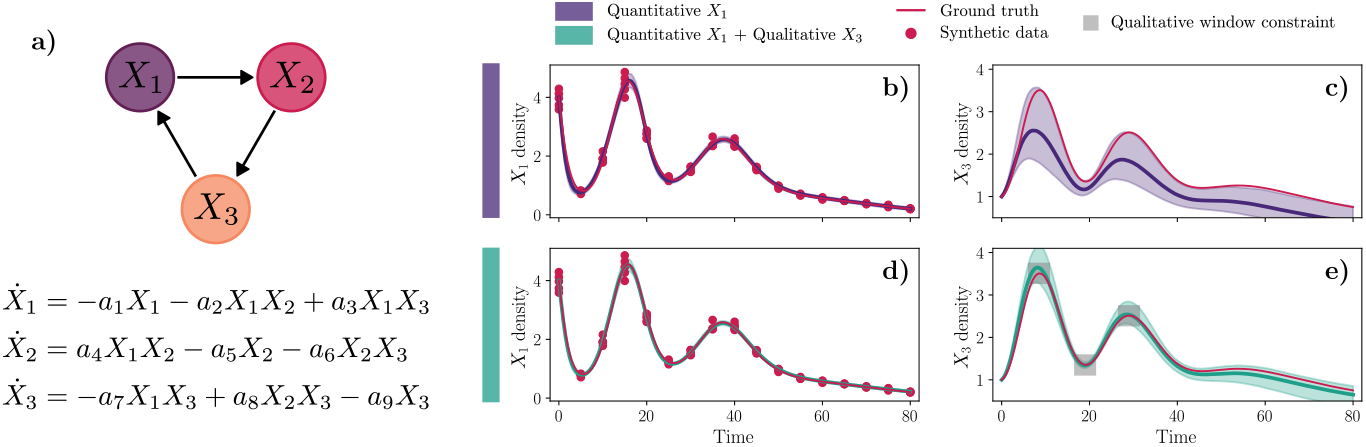
Dynamics of cycle Lotka-Volterra model for the two different approaches. A comprehensive comparison of model predictions for two approaches, classical parameter estimation and using our CrossLabFit approach. On the left is a model sketch and equations of the Lotka-Volterra model used as a testbed. In the panel of plots, each column represents the variables *X*_1_ and *X*_3_. The first row shows classical optimization results using synthetic data with log-normal noise, highlighted by red circles. The last row integrates qualitative windowing constraints for *X*_3_. The ground truth is denoted by a solid red line, contrasted with a solid line and shaded area showing the median and confidence interval of the simulations obtained by estimating parameters using nonparametric bootstrapping.

Synthetic data is generated for this 9-parameter model to evaluate the potential of the approach by specifying values for each parameter and solving the equations to simulate the dynamics. Quantitative data were derived by sampling the variable *X*_1_ at regular intervals, introducing log-normal noise at each point. For qualitative windows, we identified key peaks and valleys in the variables *X*_2_ and *X*_3_ and used the values as the center of the window, with a time width of 5 and a value height of 0.5. Using this synthetic data set, we estimated the nine parameters using a custom Differential Evolution (DE) algorithm.

On the right side of Fig 2, the plot panel shows the model results. Each column represents the model variables *X*_1_ and *X*_3_, while each row corresponds to different strategies. The Supplementary Material Fig S1 contains plots for the variable *X*_2_ and strategies related to adding qualitative windows constraints for *X*_2_.

The top row illustrates parameter fitting that involves quantitative parameter estimation using only *X*_1_ synthetic data represented by red circles. In the second row, qualitative window constraints are integrated for the variable *X*_3_. The solid red line represents the ground truth, while the solid line and shaded area illustrate the median and confidence interval of simulations resulting from the best parameter estimates obtained using our DE optimizer and nonparametric bootstrapping.

Numerical results demonstrate the impact of incorporating qualitative data into the parameter estimation process. Parameter fitting that relies solely on quantitative data serves as the baseline. Adding qualitative constraints to *X*_3_ significantly improves the fit for this variable, as shown by the tight shaded area around the solid red ground truth line in Fig 2d. In contrast, introducing qualitative constraints on *X*_2_ does not significantly improve the accuracy of the *X*_2_ simulations relative to the ground truth. Furthermore, the fourth strategy, which includes constraints on both *X*_2_ and *X*_3_, offers an improvement, but the results are similar to the third strategy, which only considers the qualitative constraint on *X*_3_, as shown in the corresponding plots in the Supplementary Material Fig S1.

These results suggest that the stochastic DE optimizer is more effective when qualitative data are integrated into the optimization process, leading to a more accurate representation of the underlying dynamics. The inclusion of qualitative constraints likely narrows the feasible parameter space, thereby accelerating the convergence of the optimization algorithm to the true parameter values.

In Fig 3a, we show violin plot illustrating the variability and distribution of the estimated parameter obtained by 1000 bootstrapping resamples using the two different data integration strategies. The width of each violin represents the density of bootstrap samples at different values, providing a visual comparison of the variability of the estimated parameter across the two strategies. The red line shows the ground truth values for each parameter. In each violin plot, the median and interquartile interval are marked as dashed lines.

**Fig 3.**
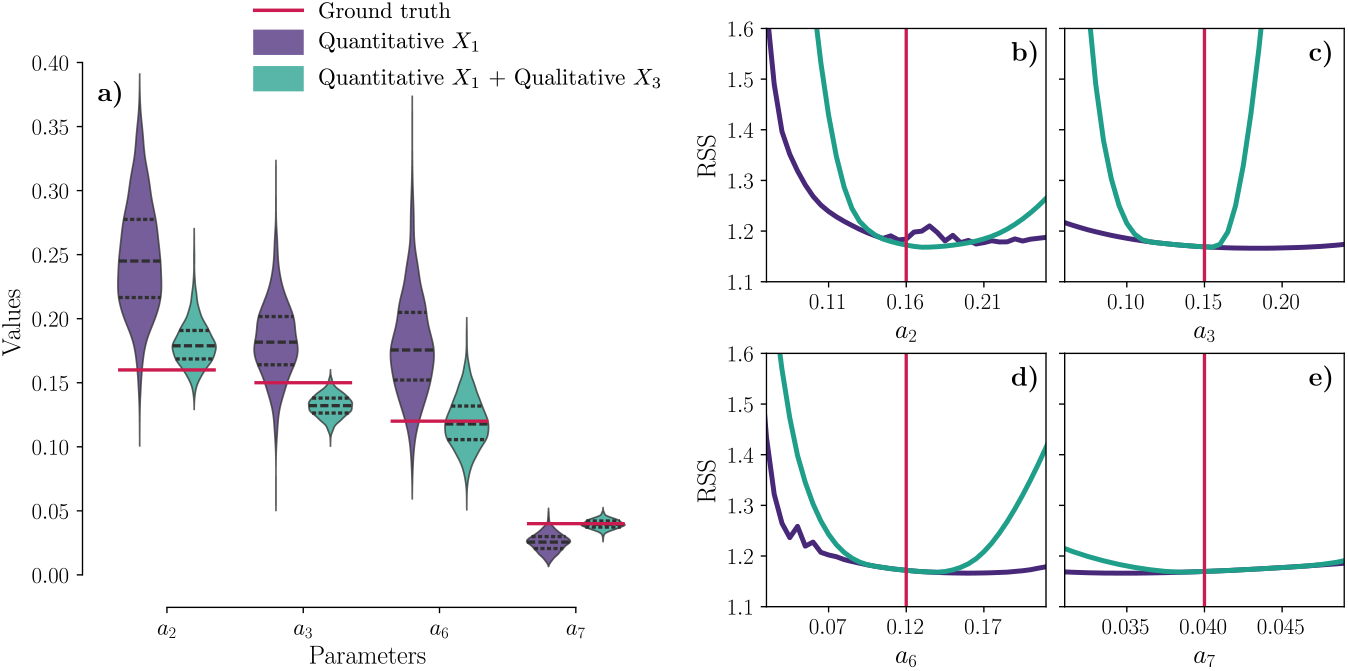
Parameter distribution and likelihood profiles for cycle Lotka-Volterra model. On the left, panel (a) shows violin plots illustrating the variability and density of the estimated parameters *a*_2_, *a*_3_, *a*_6_, and *a*_7_ derived from 1000 bootstrap resamples across the two data integration strategies. The width of each violin indicates the sample density at different values, with a red line marking the ground truth. Dashed lines within each violin represent the median and interquartile range. Panels (b-e) show likelihood profiles for four parameters in the Lotka-Volterra model (*a*_2_, *a*_3_, *a*_6_, and *a*_7_), comparing two data integration strategies (classical estimation and CrossLabFit approach) against the ground truth to assess parameter identifiability. These panels plot RSS against parameter values, indicating that minima closer to the ground truth represent more accurate estimates.

A tighter concentration of bootstrap samples around the ground truth line indicates a more accurate and consistent estimation process. The variations in the width and centering of the violins for each strategy illustrate how the inclusion of qualitative data for *X*_3_ affects the parameter estimates. The integration of qualitative constraints brings the estimates for the parameters *a*_2_, *a*_3_, *a*_6_ and *a*_7_ closer to the ground truth, with reduced variance, indicating improved estimation accuracy. In particular, the estimates for the *a*_6_ and *a*_7_ parameters are visibly improved, with the median being aligned with the ground truth values. In contrast, as observed in the Supplementary Material Fig S1, the quantitative estimation for *a*_8_ is sufficient, as the median is close to the ground truth. For the remaining parameters, the addition of qualitative information does not show any significant improvement, except for a reduction in the variance of the distributions.

Figs 3b-e shows the likelihood profiles for four parameters (*a*_2_, *a*_3_, *a*_6_ and *a*_7_) of the test-bed model. Each panel shows the RSS as a function of parameter values, with the two different strategies superimposed to illustrate their effect on parameter identifiability. The vertical red lines mark the ground truth values, which serve as a reference for the accuracy of each strategy. To obtain these curves, we chose a parameter and set its value at regular intervals throughout the search space; for each parameter value, we estimated the rest of the parameters using our DE optimizer and obtained the minimum RSS. The closer the minimum of the curves is to the ground truth, the more robust the parameter estimation is with the given strategy.

Overall, the likelihood profiles incorporating additional qualitative data are tighter than those using the classical quantitative approach, resulting in narrower confidence intervals. The profile for *a*_3_ becomes more accurate with the inclusion of *X*_3_-related qualitative data, and is closer to the ground truth. This improvement is expected given the role of *a*_3_ in the interaction between *X*_1_ and *X*_3_. The estimation of the parameters *a*_2_ and *a*_6_ also benefits from the qualitative data, showing bounds closer to the true values, while the quantitative estimation yields only a left bound. For *a*_7_, our approach shows a discernible minimum. For the remaining parameters shown in the Supplementary Material Fig S1, there is no apparent improvement, with some still showing structural non-identifiability problems. Nevertheless, the closer bounds to the ground truth suggest that our approach still offers advantages.

### Building qualitative windows from data among different labs

The last subsection shows the potential of improving parameter fitting by integrating qualitative windows. However, the main question now is how to develop these qualitative windows with experimental data from different labs. When we obtain data from different sources, we often encounter discrepancies in the values of the observables and the timing due to different measurement start points. However, we can be confident within specific ranges of values and time-frames. Fig 1b shows different examples of the types of data *e*.*g*., replicates, higher frequency measurements, confidence intervals, or even a single data point. Data sets can be mapped into a shared space where different data sets can be grouped into categories or windows. These windows can vary in size and shape and cover a sufficient range of time and values.

The primary challenge in building qualitative window constraints is comparing and merging data sets from different sources, as they may have different value scales. An approach is to normalize each dataset using its maximum and minimum values, combining all data into a common space between 0 and 1. To compare our model predictions in this normalized space, we also normalize the corresponding observables in our simulations. However, this approach preserves the shape of the observable curves of the model, regardless of their absolute values, as they are filtered by passing through the windows in the normalized space. As a result, curves of different sizes but similar shapes can be treated equivalently. To address these value scaling issues, we propose to scale the qualitative windows from the normalized space to a maximum value expected from our system.

The second challenge is to determine the shape and size of the windows. Small windows lead to more constrained trajectories in the qualitative cost function, resulting in a more limited search parameter space for the optimizer algorithm. Conversely, large windows render the qualitative constraints ineffective by allowing the entire parameter search space. The shape of the windows also affects the barriers in the parameter search space. We prefer to use square windows for computational simplicity, as it is easier to determine whether simulation values fall within a specific range than within another geometry *e*.*g*., circles. The task is to determine the number of windows and how they should span the data set. Our proposed approach involves three steps presented in the Algorithm 1, and it is more objective than manually placing windows, although manual placement is also a potential option.

#### Algorithm 1

Pipeline to build qualitative window constraints

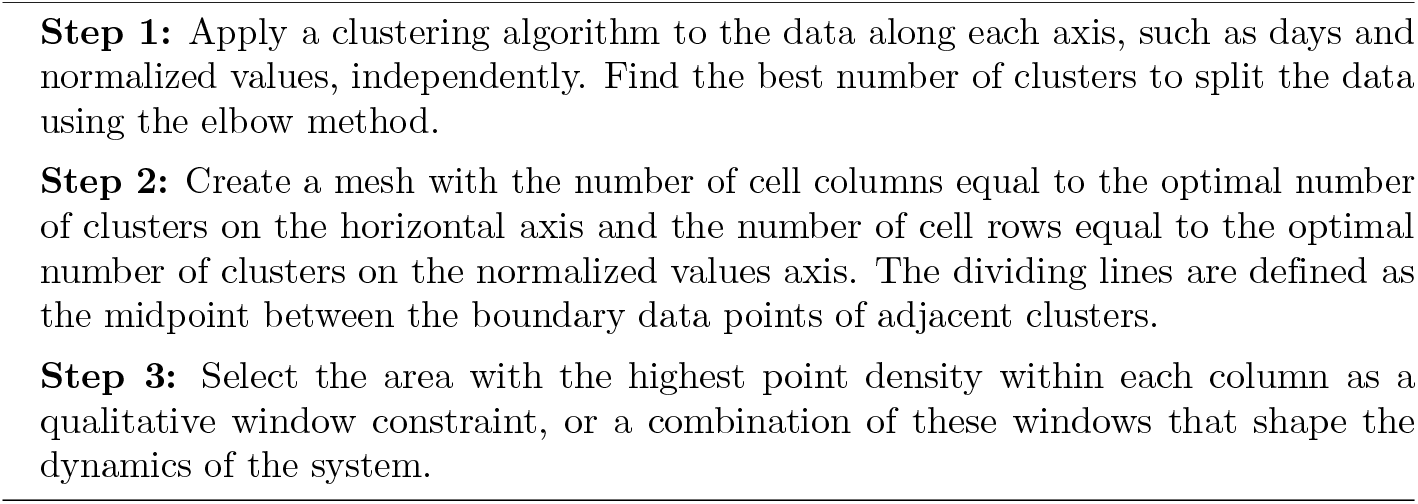

### Study case: CD8+ T cell response to influenza infection

To exemplify this process, we consider immunological responses to influenza infection, see Fig 4. We generated qualitative window constraints using CD8+ T cell data from three different sources [27–30]. We collected data from studies using consistent methods to measure CD8+ T cells in the lungs of BALB/c mice intranasally infected with influenza A PR8 (H1N1). The doses administered varied between studies, and each data set covered different frequencies and ranges of days. Each data set was normalized to its maximum and minimum values.

**Fig 4.**
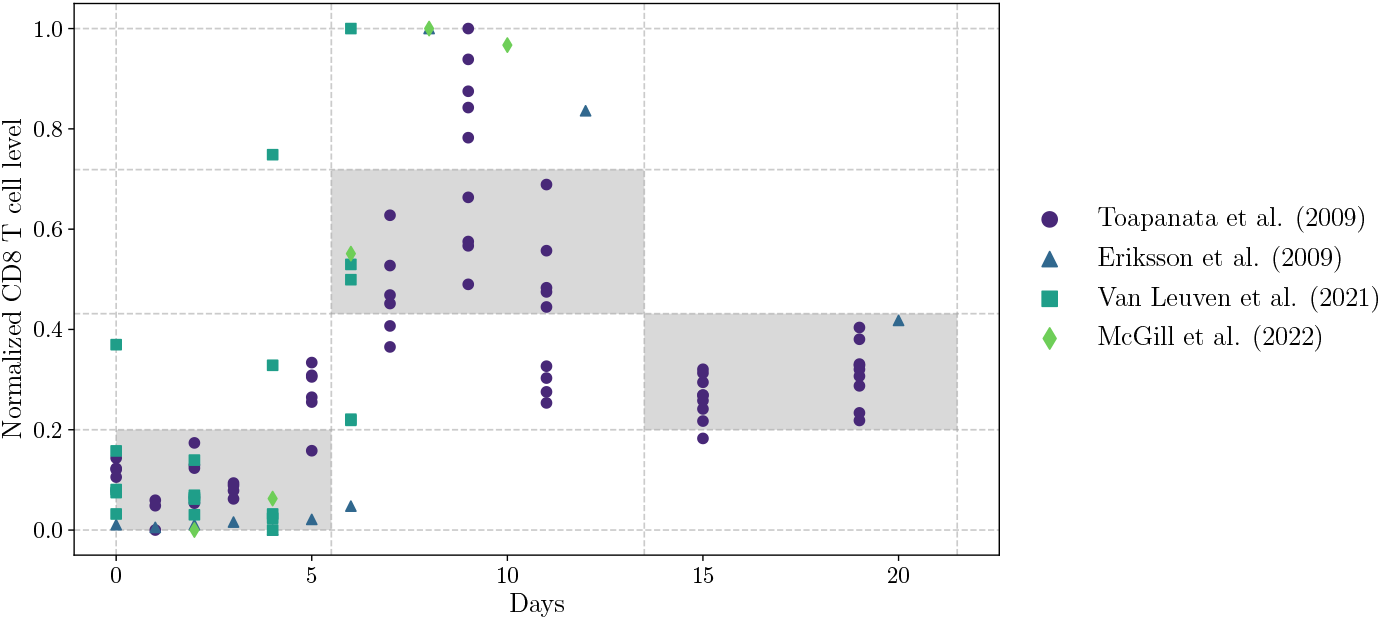
Building qualitative window constraints. The data were normalized using the minimum and maximum values of each data set. The plot shows the data from the three sources and the qualitative constraints formed by a grid where the number of vertical cells is determined by clustering the CD8+ T cell data and the number of horizontal cells is determined by clustering the days dataset. The windows were chosen according to the maximum number of points between cells in the same column; we combined two cells as the window peak value.

Fig 4 shows the four datasets and the qualitative window constraints generated from a mesh. The number of vertical cells was determined by clustering the CD8+ T cell data, while the number of horizontal cells was based on clustering the days from the datasets. We used the elbow method to determine the optimal number of clusters for both the CD8+ T cell data and the temporal aspect. The number of clusters on the temporal axis dictates the number of qualitative windows. These windows were selected based on the maximum number of data points within each column to better capture the dynamics of the CD8+ T cells.

We applied the qualitative windows from Fig 4 to a mechanistic model of Influenza A infection [31, 32]. The viral dataset from mouse lung tissues obtained from [33] serves as the quantitative data that our model aims to explain. The equations are written as follows:

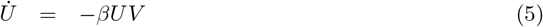

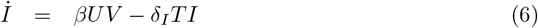

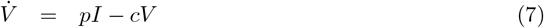

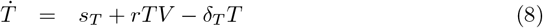

*U* represents the population of uninfected target cells, *I* denotes the population of infected cells, *V* is the concentration of virus particles, and *T* represents the population of CD8+ T cells. The parameters include the infection rate *β*, which describes how effectively the virus infects uninfected cells; the rate δ_*I*_ at which T cells kill infected cells; the replication rate *p* of new virus particles by infected cells; the clearance rate *c* of free virus particles; the proliferation rate *r* of T cells in response to the presence of the virus; and the natural death rate δ_*T*_ of T cells. The homeostatic proliferation of CD8+ T cells is *s*_*T*_ = δ_*T*_ *T*_0_, maintaining a positive level of these cells even after the virus is cleared. Here, *T*_0_ represents the initial condition of T cells, while the initial infected population is set to zero.

We performed parameter estimation using two approaches: a traditional quantitative method [34] that relies solely on the viral dataset, and our new CrossLabFit approach, which combines the viral dataset with qualitative window constraints from Fig 4. For the CrossLabFit approach, we assumed a maximum T cell level of 10^7^, scaling and shifting the qualitative windows to fit within the range of 10^6^ to 10^7^. Additionally, we imposed two constraints: first, a maximum allowable T cell level to prevent simulations from exceeding the upper limit, and second, a minimum viral load level to prevent viral dynamics from rebounding once the minimum is reached. The minimum detectable viral load is approximately 100 TCID50 (50% tissue culture infectious dose) [35]; we used 50 TCID50 as both the initial viral load and the minimum threshold for viral load constraint.

Fig 5 compares the influenza model predictions from the two approaches: classical parameter estimation and our CrossLabFit approach. In the panel of plots, each column shows the viral dynamics *V* (panels a and c) and CD8+ T cell dynamics *T* (panels b and d). The first row presents the results of classical optimization using viral load data, indicated by red circles. The second row integrates qualitative window constraints for *T* and threshold constraints for both *V* and *T*, shown as gray dashed lines. The solid lines represent the median, and the shaded areas indicate the confidence intervals of the simulations, obtained from parameter estimates using non-parametric bootstrapping.

**Fig 5.**
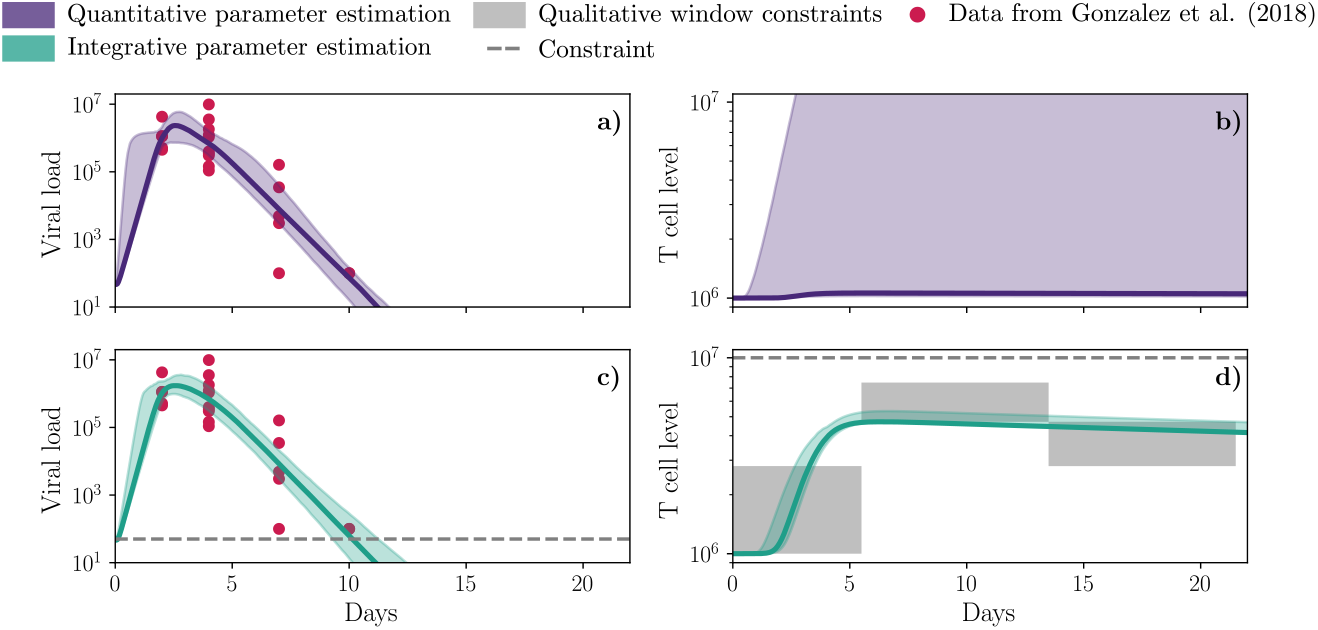
Dynamics of Influenza model for the two different approaches. A comprehensive comparison of influenza model predictions for two approaches, classical parameter estimation and using our CrossLabFit approach. In the panel of plots, each column shows the viral dynamics *V* (panels a and c) and T cell dynamic *T* (panels b and d). The first row shows classical optimization results using viral load data, highlighted by red circles. The last row integrates qualitative windowing constraints for *T*. Solid line and shaded area illustrate the median and confidence interval of the simulations obtained by estimating parameters using nonparametric bootstrapping.

Both approaches exhibit similar behavior for viral dynamics, with comparable confidence intervals. However, the CD8+ T cell dynamics differ significantly. The traditional approach shows a wide confidence interval, with the median CD8+ T cell level remaining close to the initial value. In contrast, the CrossLabFit approach produces a well-defined curve for CD8+ T cell dynamics with a much narrower confidence interval.

Fig 6a presents violin plots illustrating the variability and density of the estimated parameters *β*, δ_*I*_, *p*, and *r* derived from 1000 bootstrap resamples across the two data integration strategies. The width of each violin reflects the sample density at different values, while the dashed lines represent the median and interquartile range. Notably, the CrossLabFit approach shows a tighter distribution of values, indicating improved confidence intervals, particularly for δ_*I*_. Additionally, the parameter *r* exhibits a bimodal distribution under the quantitative estimation, in contrast to the monomodal distribution observed with the CrossLabFit approach.

**Fig 6.**
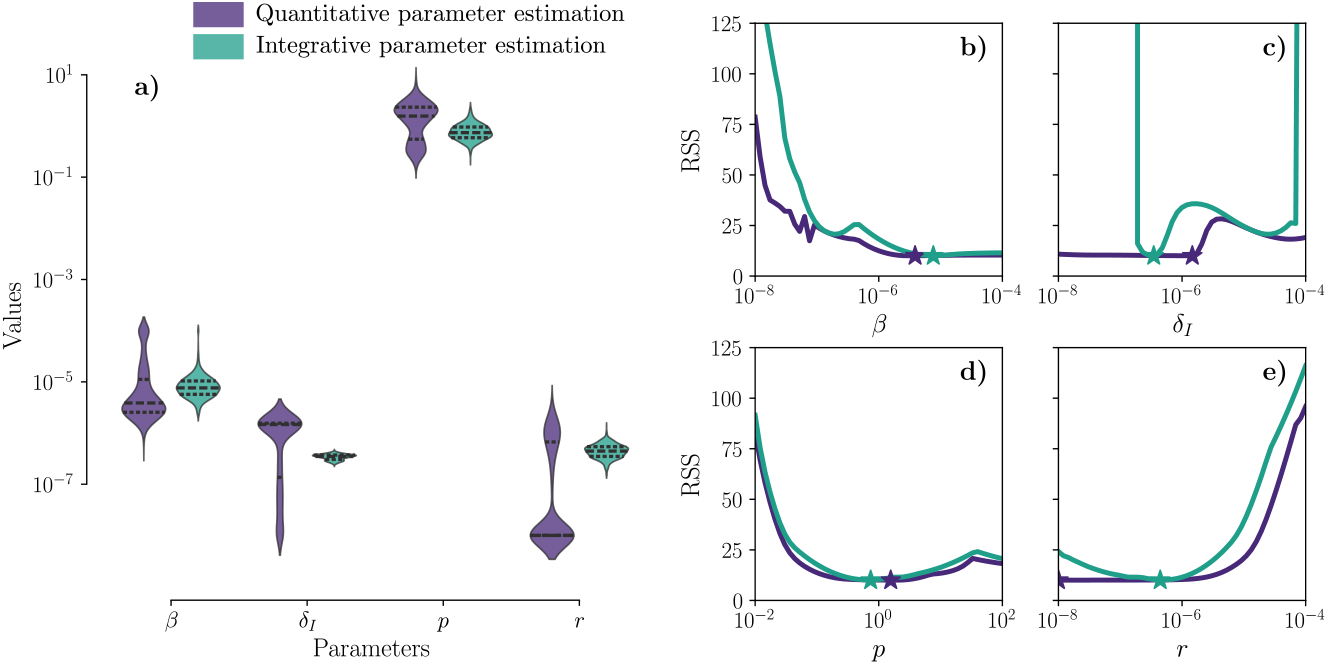
Parameter distribution and likelihood profiles for Influenza model. On the left, panel (a) shows violin plots illustrating the variability and density of the estimated parameters *β*, δ_*I*_, *p*, and *r* derived from 1000 bootstrap resamples across the two data integration strategies. The width of each violin indicates the sample density at different values. Dashed lines within each violin represent the median and interquartile range. Panels (b-e) show likelihood profiles for four parameters in the Influenza model (*β*, δ_*I*_, *p*, and *r*), comparing two data integration strategies (classical estimation and CrossLabFit approach) to assess parameter identifiability. The stars indicate the median values from the parameter distributions obtained via bootstrapping, with colors corresponding to the respective approach.

Figs 6b-e present likelihood profiles for four parameters in the Influenza model (*β*, δ_*I*_, *p*, and *r*), comparing the classical estimation with the CrossLabFit approach to assess parameter identifiability. The stars indicate the median values from the parameter distributions obtained via bootstrapping, with colors corresponding to the respective approach. For parameters *β* and *p*, there is no significant improvement in resolving the practical non-identifiability issue. While *β* shows a slightly improved left bound, *p* remains similar for both approaches. In contrast, parameters δ_*I*_ and *r* show notable improvements with the CrossLabFit approach. The likelihood profile for δ_*I*_ shifts from structural non-identifiability to having a clear minimum with tighter bounds, although an additional minimum is still observed. The parameter *r* shows a slight improvement in the left bound compared to the traditional approach. These results are expected, as δ_*I*_ and *r* are directly related to CD8+ T cell dynamics in the model equations.

Therefore, incorporating additional information in the form of qualitative windows from an unobserved variable helps identify parameters associated with that observable.

Table 1 presents the estimated parameters for the Influenza A model in mouse lungs, comparing the classical approach with the CrossLabFit approach. The values shown are the medians and 95% confidence intervals from the bootstrapped estimates displayed in Fig 6a. In addition to differences in the median values, we observe a significant improvement in the confidence intervals with the CrossLabFit approach. Notably, parameter *β* lacks an upper confidence bound in the classical approach, as the upper value reaches the limit of the search bounds. Similarly, for parameter δ_*I*_, the lower confidence bound is restricted by the lower search limit. For parameter *r*, the median lies at the lower bound of both the confidence interval and the search range in the classical approach. In contrast, the CrossLabFit approach provides well-defined confidence intervals for all parameters, also a narrower range for parameter *p* compared to the classical approach. Thus, the CrossLabFit approach enhances the ability to define confidence intervals for all estimated parameters.

**Table 1.** Comparison of estimated parameters for the Influenza A model in mouse lungs. Values are median and 95% confidence interval from bootstrapped estimates.

| Parameter | Traditional approach | CrossLabFit approach |
| --- | --- | --- |
| $\beta$ ($10^{-6}$) | 3.87 [1.45, 100] | 7.92 [3.47, 23.7] |
| $\delta_I$ ($10^{-6}$) | 1.47 [0.01, 1.71] | 0.35 [0.25, 0.41] |
| $p$ | 1.57 [0.23, 4.20] | 0.73 [0.39, 1.51] |
| $r$ ($10^{-8}$) | 1 [1, 178] | 44.2 [23.1, 78.3] |

## Discussion

The integration of qualitative data into the quantitative parameter estimation process has yielded promising results in our study. Incorporating additional information as qualitative constraints improves the search for a minimum in the cost function by excluding certain regions of the parameter space. Numerical results revealed that the CrossLabFit approach achieved a closer fit to the ground truth compared to the classical method. As shown in Fig 2, the dynamic of variable *X*_3_ was successfully replicated with the CrossLabFit approach. This suggests that incorporating qualitative constraints can effectively narrow the parameter search space, leading to more accurate model predictions and improving identifiability analysis. Such findings align with previous research that emphasizes the value of qualitative information in complex system modeling [14, 20, 21].

Bootstrapping analysis in Fig 3a revealed that the variability of parameter estimates generally decreased when qualitative data were included. However, the accuracy of the estimated values is improved only for the parameters *a*_6_ and *a*_7_, since the medians hit the ground truth value. This enhancement in the precision of parameter estimates is particularly notable for parameters that directly interact with the variable *X*_3_ for which qualitative data were generated. Likelihood profiles in Figs 3b-e also indicated that some parameters associated with qualitative data constraints had tighter likelihood profiles, demonstrating the improvement of identifiability.

However, our approach has limitations. Our proposed approach uses a hard-penalty method, where violations of qualitative windows impose infinite penalties. This assumes confidence in the placement of qualitative windows. Hard penalties can exclude valid parameter sets and reduce robustness. Alternative approaches will be investigated, such as soft constraints that proportionally penalize deviations.

Our approach does not improve identifiability for all parameters *e*.*g*., *a*_5_, suggesting potential structural non-identifiability issues that our approach may not resolve.

Parameter *a*_5_ is related to the deletion rate of the variable *X*_2_, and in general, adding qualitative windows to this variable does not improve the estimation of the parameters.

Supplementary Material Fig S2 and Fig S3 include results for two additional test bed models. For the linear-chain Lotka-Volterra model without a feedback loop, parameter estimation enhancements were predominantly observed for the qualitative constraints on *X*_3_. Conversely, in the modified Lotka-Volterra model where *X*_2_ and *X*_3_ interact solely with *X*_1_ and not with each other, the integration of qualitative data for both *X*_2_ and *X*_3_ yielded improvements in parameter estimation.

In our study, the algorithm to build qualitative windows plays a crucial role in integrating diverse data sources into the parameter estimation process. We chose to normalize each dataset independently, as the varying scales between datasets posed a challenge. Specifically, larger datasets tended to overshadow smaller ones, making it difficult to reflect the dynamics of the smaller datasets in the model. By normalizing each dataset to its minimum and maximum values, we ensured that all data sources were given equal weight in the qualitative window construction. In Fig 4, we can observe all the normalized data using this approach. When we attempted to normalize the datasets as a group, however, we found that the data from [27, 28] had larger values, which flattened the dynamics of the other datasets. Alternative standardization techniques could be explored to refine this process further.

We applied our approach to influenza infection in the lungs of mice. Bootstrapping results showed significant improvement for T cell dynamics, as illustrated in Fig 5.

When parameter estimation relied solely on viral load, the CD8+ T cell dynamics remained nearly flat.

Our findings underscore the selective utility of integrating data from different labs into qualitative data for constructing more complex models. Numerical results on test bed models highlight that the key to improving parameter estimation lies not only in the quantity of information but also in the time position of the qualitative window and the strategic selection of variables for qualitative integration. This pioneering approach will lead to significant improvements in computational biology.

## Materials and methods

### Test bed model and synthetic data

The model shown in Fig 2 is one of three Lotka-Volterra models, each incorporating different interactions between the observable variables *X*_1_, *X*_2_, and *X*_3_. The parameters *a*_*i*_ for *i* ∈{0, …, 9} define the different models. In the model shown in Fig 2, the input term *a*_0_*X*_1_ was omitted from the rate of change equation for 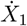 because its value was set to zero. For each model, we selected arbitrary parameter values designed to produce damped oscillations with peaks and valleys; a complete list of parameters is provided in Table 2.

**Table 2.** Parameters for the three Lotka-Volterra models used to generate synthetic data.

| Parameter | Linear | Cycle | 2-predators |
| --- | --- | --- | --- |
| $a_0$ | 0.14 | 0.00 | 0.45 |
| $a_1$ | 0.00 | 0.18 | 0.00 |
| $a_2$ | 0.16 | 0.16 | 0.16 |
| $a_3$ | 0.00 | 0.15 | -0.15 |
| $a_4$ | 0.15 | 0.11 | 0.15 |
| $a_5$ | 0.11 | 0.02 | 0.11 |
| $a_6$ | 0.05 | 0.12 | 0.00 |
| $a_7$ | 0.00 | 0.04 | -0.08 |
| $a_8$ | 0.06 | 0.12 | 0.00 |
| $a_9$ | 0.05 | 0.02 | 0.05 |

Synthetic data for *X*_1_ were generated using these parameter values by solving the ODEs with the odeint function from the SciPy library. We sampled the resulting dynamics of each observable every 5 time units. For each sampled point, we generated 5 random samples using a log-normal distribution. This approach was chosen for two reasons: first, it introduces greater variation at higher values, which is typical of real biological data; and second, it avoids generating negative values, which would not make sense in this system. The qualitative windows were centered around the peaks and valleys of *X*_2_ and *X*_3_. The time range (width) of each window was fixed at 5 units, and the concentration range (height) was set at 0.5. This synthetic data and qualitative constraints were provided as input to our custom-built DE optimizer. All data generation, analysis, and plotting was done in Python, while parameter estimation and bootstrapping was done for our custom DE optimizer coded in CUDA/C.

### Qualitative windows constraints

To build the qualitative window constraints, we used raw datasets from [27, 30] and extracted data from plots in [28, 29] using the PlotDigitizer web app [36]. Each dataset was normalized independently using its respective maximum and minimum values, after which all datasets were combined into one. We employed the `KMeans` clustering function from the `Scikit-learn` library with default parameters. The clustering was performed separately for the time data and the normalized values, varying the number of clusters for each to determine the optimal count using the elbow method. Once the optimal number of clusters was identified for both the time and normalized variable values, we followed the steps outlined in Algorithm box 1 to define the qualitative window constraints. The boundaries of these windows were saved in a file and provided as input to our custom-built DE optimizer. All processes were carried out in Python.

### High-Performance Computing

The code for bootstrapping and identifiability analysis was implemented in CUDA/C to leverage high-performance computing on the Falcon supercomputer [37], utilizing GPU nodes. Falcon is located at the Idaho National Laboratory Collaborative Computing Center (C3) in Idaho Falls, Idaho, USA. We conducted 1000 simulations for each case study and 50 simulations for each likelihood profile. Each optimization run took approximately 20 minutes.

### GPU-accelerated version of Differential Evolution Optimizer

To optimize the cost function, we employed a custom-built Differential Evolution (DE) algorithm [38], chosen for its simplicity and effectiveness in various applications [39].

We developed a GPU-accelerated version of the DE algorithm using CUDA/C, significantly enhancing computational speed by leveraging GPU parallelization, following the approaches of previous studies [40, 41].

Additionally, we implemented a fifth-order Runge-Kutta method to solve the ODE system within our optimizer. Within the DE algorithm, we utilized a population array consisting of 8192 parameter sets and 10000 iterations for the main loop. The initial population was randomly sampled from a uniform distribution spanning the minimum and maximum values of each parameter. To generate a new mutant vector within the population, we adopted the DE/rand/1/bin strategy. This approach involves selecting three random vectors (sets of parameters) from the population and using them to mutate each vector within the population. The mutation was performed with a mutation factor of *F*_*m*_ = 0.8, a crossover rate of *C*_*r*_ = 0.8, and a binomial criterion selection.

A detailed description of the algorithm is provided in Supplementary Material Algorithm S1.

### Bootstraping

We applied a nonparametric bootstrap approach using Monte Carlo resampling. Data were resampled with replacement to create a sample of the same size as the original dataset. Parameters were then estimated from each resampled dataset. For bootstrapping [42], we resample the synthetic data with replacement and then estimate the parameters from the resample. We run 1000 bootstrap parameter estimates for each strategy. This allowed us to obtain the corresponding parameter distributions by refitting the model in each of these iterations.

### Identifiability analysis

A model is considered identifiable when its parameters can be uniquely determined. We used the profile likelihood method [4], where each parameter is varied while the others are re-optimized. This approach detects both structural and practical non-identifiability [4, 43]. Structural non-identifiability arises from the model structure, while practical non-identifiability is due to data quality or quantity. A parameter is identifiable if its profile likelihood is concave; a flat valley indicates practical non-identifiability, and a constant cost function suggests structural non-identifiability.

## Data and Code availability

The codes and data generated and/or analyzed during the current study are available in the GitHub repository https://github.com/systemsmedicine/CrossLabFit

## Acknowledgments

Research reported in this publication was supported by the National Institute of General Medical Sciences of the National Institutes of Health under award numbers R01GM152736 and P20GM104420. The content is solely the responsibility of the authors and does not necessarily represent the official views of the National Institutes of Health.

## Supporting information

**Fig S1 Results for the cyclic Lotka-Volterra model**. The panel displays a sketch of the model, the equations used, simulation results for each strategy, parameter distributions, and likelihood profiles comparing the strategies. The strategies involve parameter estimation using synthetic data from *X*_1_ alone, with qualitative constraints in *X*_2_, in *X*_3_, and in both *X*_2_ and *X*_3_. All plots are color-coded according to the key labels above for consistency across strategies.

**Fig S2 Results for the linear Lotka-Volterra model**. The panel displays a sketch of the model, the equations used, simulation results for each strategy, parameter distributions, and likelihood profiles comparing the strategies. The strategies involve parameter estimation using synthetic data from *X*_1_ alone, with qualitative constraints in *X*_2_, in *X*_3_, and in both *X*_2_ and *X*_3_. All plots are color-coded according to the key labels above for consistency across strategies.

**Fig S3 Results for the 2-predator Lotka-Volterra model**. The panel displays a sketch of the model, the equations used, simulation results for each strategy, parameter distributions, and likelihood profiles comparing the strategies. The strategies involve parameter estimation using synthetic data from *X*_1_ alone, with qualitative constraints in *X*_2_, in *X*_3_, and in both *X*_2_ and *X*_3_. All plots are color-coded according to the key labels above for consistency across strategies.

**Algorithm S1 Pseudocode for a GPU-accelerated Differential Evolution algorithm**. The algorithm shows the pseudocode for a GPU-accelerated Differential Evolution algorithm with qualitative constraints for parameter estimation.

