## Supplementary for "CrossLabFit: A Novel Framework for Integrating Qualitative and Quantitative Data Across Multiple Labs for Model Calibration"

### Cyclic Lotka-Volterra model:

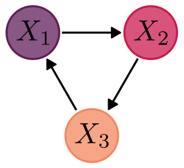


$$\dot{X}_1 = -a_1 X_1 - a_2 X_1 X_2 + a_3 X_1 X_3$$
$$\dot{X}_2 = a_4 X_1 X_2 - a_5 X_2 - a_6 X_2 X_3$$
$$\dot{X}_3 = -a_7 X_1 X_3 + a_8 X_2 X_3 - a_9 X_3$$

#### Simulations:

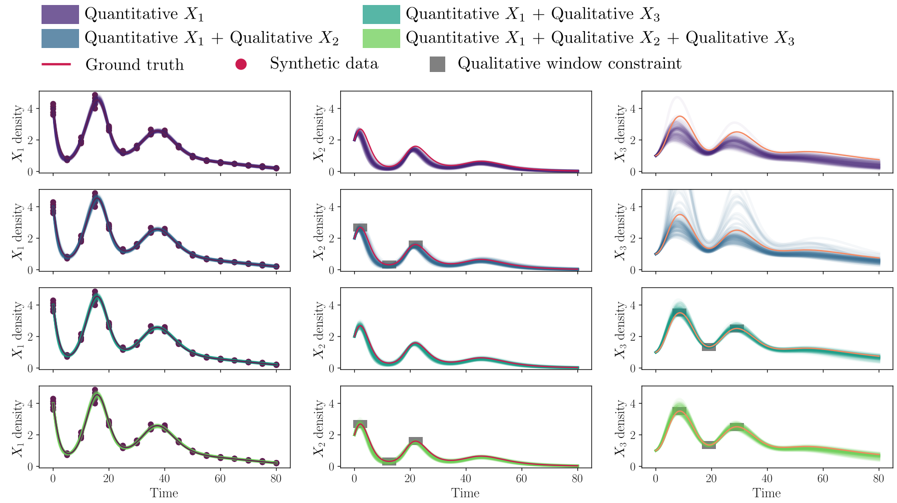


#### Parameter distributions:

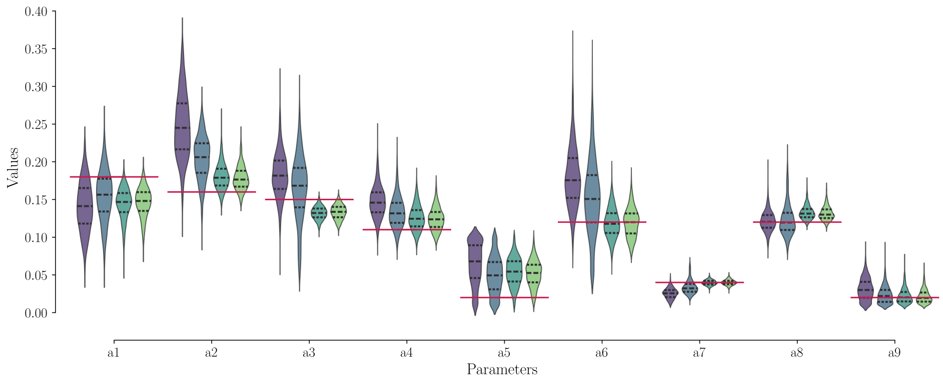


#### Likelihood Profiles:

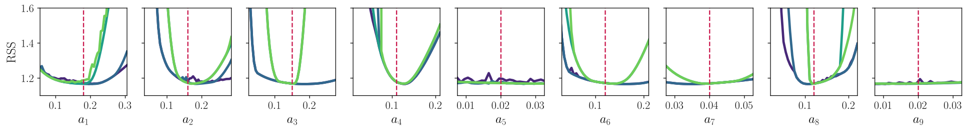


**Fig S1.** Results for the cyclic Lotka-Volterra model. The panel displays a sketch of the model, the equations used, simulation results for each strategy, parameter distributions, and likelihood profiles comparing the strategies. The strategies involve parameter estimation using synthetic data from $X_1$ alone, with qualitative constraints in $X_2$, in $X_3$, and in both $X_2$ and $X_3$. All plots are color-coded according to the key labels above for consistency across strategies.

### Linear Lotka-Volterra model:

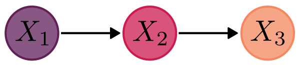


$$\dot{X}_1 = a_0 X_1 - a_2 X_1 X_2$$
$$\dot{X}_2 = a_4 X_1 X_2 - a_5 X_2 - a_6 X_2 X_3$$
$$\dot{X}_3 = a_8 X_2 X_3 - a_9 X_3$$

#### Simulations:

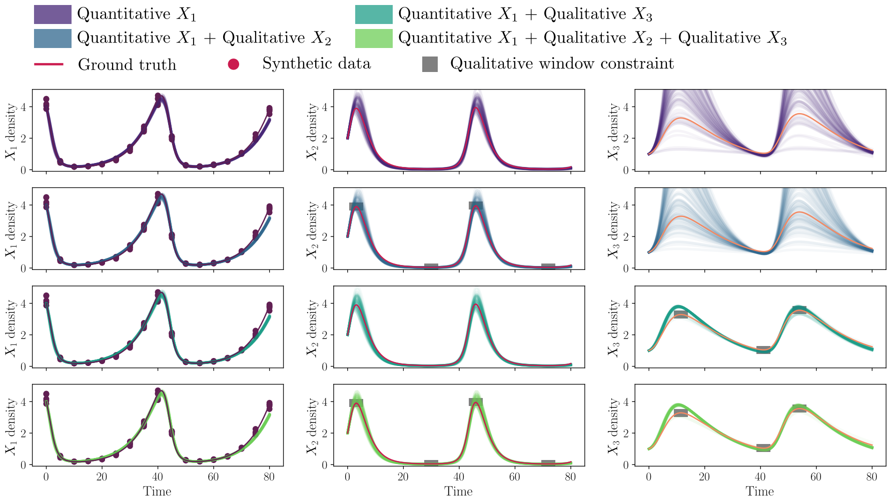


#### Parameter distributions:

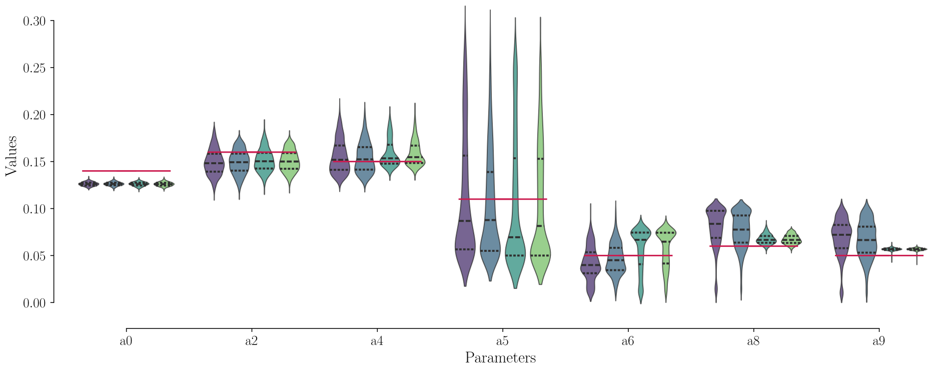


#### Likelihood Profiles:

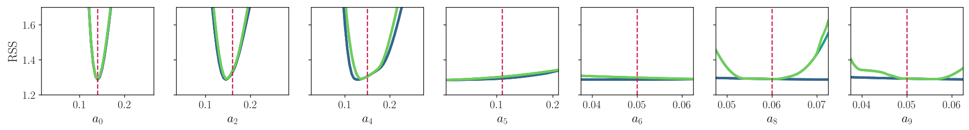


**Fig S2.** Results for the linear Lotka-Volterra model. The panel displays a sketch of the model, the equations used, simulation results for each strategy, parameter distributions, and likelihood profiles comparing the strategies. The strategies involve parameter estimation using synthetic data from $X_1$ alone, with qualitative constraints in $X_2$, in $X_3$, and in both $X_2$ and $X_3$. All plots are color-coded according to the key labels above for consistency across strategies.

### 2-predator Lotka-Volterra model:

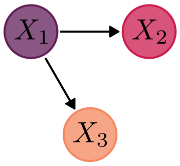


$$\dot{X}_1 = a_0 X_1 - a_2 X_1 X_2 - a_3 X_1 X_3$$
$$\dot{X}_2 = a_4 X_1 X_2 - a_5 X_2$$
$$\dot{X}_3 = a_7 X_1 X_3 - a_9 X_3$$

#### Simulations:

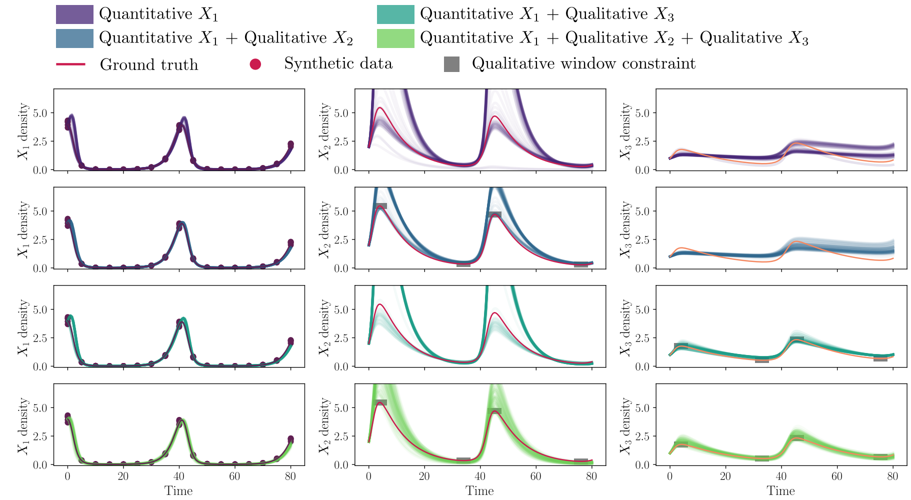


#### Parameter distributions:

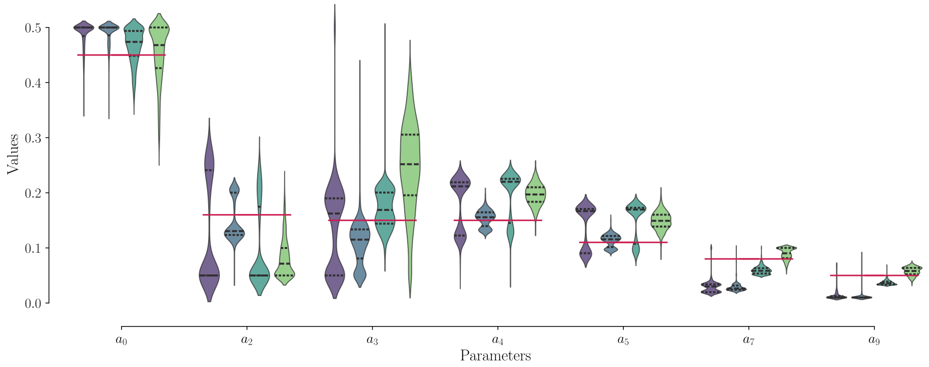


#### Likelihood Profiles:

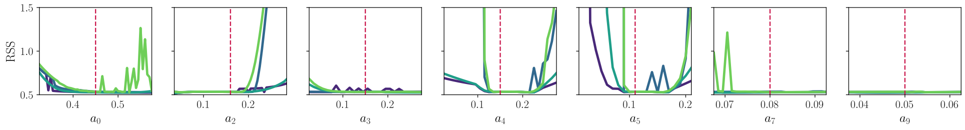


**Fig S3.** Results for the 2-predator Lotka-Volterra model. The panel displays a sketch of the model, the equations used, simulation results for each strategy, parameter distributions, and likelihood profiles comparing the strategies. The strategies involve parameter estimation using synthetic data from $X_1$ alone, with qualitative constraints in $X_2$, in $X_3$, and in both $X_2$ and $X_3$. All plots are color-coded according to the key labels above for consistency across strategies.

**Algorithm S1** Pseudocode for a GPU-accelerated Differential Evolution algorithm with qualitative constraints for parameter estimation.

**Input:** $N$: population size, $C_r$: crossover probability, $F_m$: mutation factor, $D$: Number of parameters to estimate, $M$: penalty factor.

**function** COSTFUNCTION(Pars[$i$], Data$_{qt}$, $\Delta t_{\text{ql}}$, $\Delta X_{\text{ql}}$)
  Data$_{\text{sim}}$ $\leftarrow$ 5THRUNGE-KUTTASOLVER(ODEmodel, Pars[$i$])
  **if** Data$_{\text{sim}}$ touches all qualitative windows defined for each square $\Delta t_{\text{ql}} \times \Delta X_{\text{ql}}$ **then**
    **return** $\sum_n$ (Data$_{\text{sim}}[n]-$ Data$_{\text{qt}}[n])^2$
  **else**
    **return** $M$
  **end if**
**end function**

**Initialization:** Generate a uniform random population vector Pars consisting of $N$ arrays, each containing $D$ parameters to estimate. Then, evaluate the cost function using the quantitative data Data$_{\text{qt}}$ and qualitative boundary constraints, defined by $\Delta t_{\text{ql}}$ (window time width) and $\Delta X_{\text{ql}}$ (window variable value height).
**parfor** $i \leftarrow 0$ to $N$ **do**
  **for** $j \leftarrow 0$ to $D$ **do**
    Pars[$iD+j$] $\leftarrow$ uniform random value within the search space for the parameter $j$
  **end for**
  J[$i$] $\leftarrow$ COSTFUNCTION(Pars[$i$], Data$_{\text{qt}}$, $\Delta t_{\text{ql}}$, $\Delta X_{\text{ql}}$)
**end parfor**

**Optimization:** Main loop for optimization using the DE method with the mutation strategy DE/rand/1/bin.
**for** iterations $\leftarrow$ 0 to maximum iterations **do**
  **parfor** $i \leftarrow 0$ to $N$ **do** ▷ Each thread $i$ of the GPU perform the code below
    $j_{\text{rand}} \leftarrow \text{random}(0, D)$
    **for** $j \leftarrow 0$ to $D$ **do** ▷ Generate a new mutated population
      **if** $\text{random}(0,1) < C_r$ **or** $j$ is $j_{\text{rand}}$ **then**
        $x, y, z \leftarrow \text{random}(0, N)$ **and** $x \neq y \neq z \neq i$
        newPars[$iD+j$] $\leftarrow$ Pars[$xD+j$] $+ F_m$(Pars[$yD+j$] $-$ Pars[$zD+j$])
      **else**
        newPars[$iD+j$] $\leftarrow$ Pars[$iD+j$]
      **end if**
    **end for**
    $J_{\text{new}}[i] \leftarrow$ COSTFUNCTION(newPars[$i$], Data$_{\text{qt}}$, $\Delta t_{\text{ql}}$, $\Delta X_{\text{ql}}$) ▷ Evaluate the new population
    **if** $J_{\text{new}}[i] < J[i]$ **then** ▷ Selection of a better parameter set
      $J[i] \leftarrow J_{\text{new}}[i]$
      **for** $j \leftarrow 0$ to $D$ **do** Pars[$iD+j$] $\leftarrow$ newPars[$iD+j$]
      **end for**
    **end if**
  **end parfor**
**end for**

**Finalization:** Searches for the index $i_{\text{min}}$ with the lower value in the vector $J$ and returns the best estimate of the parameters in the population Pars[$i_{\text{min}}D+j$] where $j \leftarrow 0$ to $D$.
